# Serine Recombinase PinR Inverts Cryptic Prophage DNA to Block Adsorption of Phages

**DOI:** 10.1101/2025.03.28.646004

**Authors:** Joy Kirigo, Daniel Huelgas-Méndez, Michael J Benedik, Rodolfo García-Contreras, Thomas K. Wood

## Abstract

Recombinases catalyze site-specific integration, excision, and inversion of DNA and are found in myriad defense islands; however, their function in phage-defense is unknown as they are frequently dismissed as markers of prophages. Here, we characterize the physiological role of the previously-uncharacterized serine recombinase PinR of *Escherichia coli* cryptic prophage *rac* and discover that it inhibits T2 phage infection by inverting a 1,797 bp segment in a different cryptic prophage *e14* to inhibit T2 infection; this inversion leads to the formation of a novel protein from two spliced genes, StfE2, that we find blocks phage adsorption. Modeling shows StfE2 inhibits T2 phage adsorption by preventing Gp38 binding to its primary receptors porins FadL and OmpF. Corroborating the receptor-blocking hypothesis, T2 escape mutants evolve resistance to PinR phage defense by mutating *gp38* to remove 16 aa in the hyper variable region 3. Therefore, we discovered the first recombinase-activated phage inhibition system.

## INTRODUCTION

The symbiotic relationship between prophages (integrated viral genomes) and their hosts is longstanding and widespread (1). The majority of all sequenced bacterial genomes contain one or more prophage sequences (1,2), and these pervasive viral genomes are present not only in bacterial genomes (3) but also plant (4), animal, and human genomes (5). Active prophages provide reservoirs for virulence genes (e.g., Shiga toxin, botulinum toxin, diphtheria toxin) (6), and host immunity against phages (e.g., restriction systems) (7).

In contrast, cryptic prophages; i.e., prophage fossils that are no longer able to lyse the cell or make phage particles (8), have only recently been shown to play a role in bacterial physiology. For example, *Escherichia coli* K-12 contains nine cryptic prophages (*CP4-6, DLP12, e14, rac, Qin, CP4-44, CPS-53, CPZ-55* and *CP4-57*) (9), and they have been shown to influence its stress response to antibiotics and oxidative stress (10,11) as well as regulate resuscitation from stress-induced dormancy (12). Moreover, like prophages, cryptic prophages are involved in phage defense in that *e14* prophage encodes phage inhibition systems Lit (inhibits T4 replication) (13) and McrA (degrades T-even phage DNA) (14), and *CP4-6, rac, Qin, CP4-44,* and *CP4-57* contain toxin-antitoxin systems (9), whose primary physiological role is phage inhibition (15,16).

Cryptic prophages are also hotbeds for DNA-modifying enzymes, with site specific recombinases (integrases) that remain active and are critical for innovations in biotechnology (17) and synthetic biology (18). *CP4-6, CP4-57, CPS-53, CPZ-55, DLP12, e14, rac* and *Qin* prophages all contain recombinases (9), which catalyze the integration, excision, or inversion of defined DNA segments (19). These recombinases fall into two main families, serine recombinases and tyrosine recombinases, named in reference to the conserved amino acid residue that mediates catalysis (19). Unlike tyrosine recombinases, without other factors (e.g., recombination directionality factor) (20), serine recombinases catalyze only irreversible recombination, which makes them powerful genetic tools (18). Serine recombinases fall into two classes, large serine recombinases, responsible for excision, integration, deletion, or transposition, and small serine recombinases (SSR) that function as invertases or resolvases (21).

Serine recombinases also play an important role in host and phage physiology. For example, SSR Hin catalyzes DNA inversion of a 1 kb segment that controls flagella phage variation in *Salmonella typhimurium* (22), increasing bacterial virulence (23). Gin and Cin (SSRs) catalyze inversions of the 3 kb (G segment) and 4.2 kb (C segment) of phages Mu (24) and P1 (25), respectively. Inversion in both the G- and C- segments enables the phages to adsorb to different bacterial hosts (24,26). *E. coli* K-12 also contains a SSR in cryptic prophage *e14*, PinE, which inverts a 1.8 kb segment containing putative tail fiber genes; however, the function of this inversion is not clear (27).

Previously, while determining that some *E. coli* cells that survive phage infection are dormant, we found point mutations arise in *pinR* and *pinQ*, which encode uncharacterized putative SSRs, as well as found deleting *pinR* increases the sensitivity to T2 phage 330-fold (28). Hence, we sought to characterize the physiological role of PinR here (PinQ is 99% homologous so it was not studied further). Critically, this is the first report of a recombinase playing a role in phage defense even though recombinases are common in phage-defense islands, but they have been dismissed (29). We found that upon T2 infection, PinR inverts a 1.8 kb segment in *e14*; this inversion inhibits T2 phage by creating a new, spliced protein encoded by the upstream portion of *stfE* and from the complementary strand of *stfP* after the inversion. StfE2 is found to reduce adsorption of phage T2 and modeling shows StfE2 likely inhibits T2 by blocking access to its *E. coli* receptor, FadL. Moreover, T2 escapes the PinR-mediated inversion through mutation of *gp38,* which encodes its adhesion protein. Therefore, we discovered a new class of phage inhibition system based on DNA recombination and a spliced protein.

## MATERIALS AND METHODS

### Bacterial strains, plasmids, phages, medium, and antibiotics

The bacterial strains (*E. coli* BW25113 (30) and its isogenic mutants), plasmids, and phages used in this work are described in **Table S1**. Cells were grown in lysogeny broth (LB, 1 % tryptone, 0.5 % yeast, and 1 % NaCl w/v) at 37°C. Kanamycin, 50 µg/mL (Kan50) was used for preculturing knockout mutants, and chloramphenicol, 30 µg/mL (Cm30), was added to strains containing pCA24N-based plasmids to maintain the vector. Single-gene knockouts of *E. coli* were obtained from the Keio Collection (30). Plasmids used in this study, except for pCA24N-*stfE2* and pBS(Kan)-*stfP2*, were obtained from the ASKA collection (31) and gene expression was induced by the addition of isopropyl β-*D*-1-thiogalactopyranoside (IPTG, 1 mM). pCA24N-*pinR* was cured from BW25113 cells isolated from T2 plaques by growing in overnight LB liquid cultures with IPTG in the absence antibiotic for the plasmid and by screening using LB and LB chloramphenicol plates. The absence of the plasmid was confirmed via PCR using pCA24N-specific primers (pCA24N_Fow and pCA24N_Rev, **Table S2**). Phage lysates were stored at 4°C, and T2 phage was used at a multiplicity of infection (MOI) ∼ 0.01 for all experiments unless stated otherwise.

pCA24N-*stfE2* was constructed by amplifying a 422 bp fragment from the inverted region of the chromosome of PinR-producing cells (BW25113/pCA24N-*pinR*) using primers StfE2_Fow and StfE2_Rev (**Table S2**) to introduce restriction sites for StuI and PstI (31). pBS(Kan)-stfP2 was constructed by amplifying an 833 bp fragment from the inverted region of the chromosome of PinR-producing cells (BW25113/pCA24N-*pinR*) using primers StfP2_Fow and StfP2_Rev (**Table S2**) to introduce restriction sites for ApaI and BamHI to facilitate cloning into pBS(Kan) (32); pBS(Kan) was used since it has better promoter silencing with glucose than pCA24N and construction in pCA24N led only to mutations in *stfP2*. Constructed plasmid sequences were verified by Plasmidsaurus.

### Temporal turbidity and cell viability

T2 phage was added to exponentially-growing cells (turbidity at 600 nm ∼ 0.5), and the cells were incubated at 37°C and 250 rpm and monitored using a UV-Vis spectrometer. For cell viability, 1 mL aliquots were washed twice with phosphate-buffered saline (PBS, 8 g NaC1, 0.2 g KC1, 1.15 g Na_2_HPO_4_, and 0.2 g KH_2_PO_4_ in ddH_2_O 1000 mL), and enumerated using the drop assay (33).

### Phage titers and plaque assay

For phage titers, 1 mL samples were centrifuged (10 min, 5,000 rpm) to separate free phage from cells, and 100 µL samples of the supernatant (containing free phage) were serially diluted in phage buffer (0.1 M NaCl, 10 mM MgSO_4_, and 20 mM Tris-HCl pH 7.5). Plaques formed (PFU/mL) were enumerated on double-layer agar plates (1% tryptone, 0.5% NaCl, 1.5% agar lower layer and 0.4% agar top layer) using the drop assay.

### Sequencing genomes of cells producing PinR that survive T2 infection

To gain insights into the mechanism of PinR phage inhibition, PinR was produced for 16 h via BW25113/pCA24N-*pinR*. To double layer agar plates, 100 µL of each overnight culture and 100 µL drop of T2 phage stock (6 x 10^8^ PFU/mL) were added and incubated overnight. Surviving colonies inside the plaque were purified from phages by streaking on LB Cm30 plates. Chromosomal DNA was extracted using the Qiagen DNeasy Ultraclean Microbial Kit then sequenced using Oxford Nanopore Technology and genome assembled using Flye v2.9.1 by Plasmidsaurus after removing the sequence reads of T2 DNA. Accession numbers for all the sequences are shown in **Table S3**.

### DNA inversion via quantitative polymerase chain reaction (qPCR)

qPCR was used to quantify the amount of inverted chromosomal DNA in overnight cultures containing pCA24N-derived plasmids that were induced with IPTG. Genomic DNA was extracted using the Qiagen DNeasy Ultraclean Microbial Kit. Primers (**Table S2**) were used to amplify a noninverted 153 bp (PinR_1 and PinR_2) or noninverted 831 bp (PinR_1B and PinR_2B) or inverted 494 bp (PinR_1 and PinR_3) or inverted 586 bp (PinR_1B and PinR_3B) segments of the *E. coli* chromosome of the *e14* cryptic prophage (cDNA coordinates: 1,196,443 – 1,210,635) using the Luna Universal qPCR Master Mix. (NEB #M3003). No housekeeping primers are required since inversion compares the ratio of expression of inverted versus noninverted in the same DNA sample.

### T2 adsorption

Phage adsorption (34,35) was assayed by diluting overnight cultures 1:100 into 10 mL of LB (or LB Cm30 for pCA24N-based strains) and incubated until turbidity (600 nm) ∼ 0.5, then T2 phage was added to the cultures, which were incubated at room temperature. Cells were subsequently removed by centrifugation, and samples (1 mL) were taken after 0, 8, and 16 min, and used to quantify the remaining phages not adsorbed to the cells using the plaque assay. As a complementary method, phages attached after two washes with PBS (5,000 rpm for 10 min) to remove free and loosely-attached phage were enumerated after resuspending in 1 mL PBS and using chloroform to release attached phages using the plaque assay.

### Antibiotic import through FadL and OmpF

To determine the extent of antibiotic import through the FadL and OmpF pores after production of PinR, ampicillin (100 µg/mL, 10 minimum inhibitory concentration, MIC) or ciprofloxacin (5 µg/mL, 10 MIC) were added to exponentially-growing cells after 2 h production of PinR via pCA24N-*pinR*, and the cells were cultured for 3 h. After treatment, cultures were washed twice with PBS and cell viability (colony forming units, CFU/mL) was determined using the drop assay. Temporal turbidity (600 nm) measurements were also determined over 10 h every 10 min using a Tecan Sunrise UV-Vis spectrometer for overnight cultures diluted 1:100 into fresh LB in 96-well plates that were incubated at 37℃ with shaking (250 rpm) until the exponential growth phase (turbidity ∼ 0.3), followed by the addition of ampicillin at varying concentrations (0, 5, 10, 20, 50 µg/mL).

### Molecular docking

Protenix (36) was used to predict protein-protein interactions. Structures for protein docking were predicted with AlphaFold3 (37). 3D-Structures of Gp38, FadL and OmpF were downloaded from the Protein Data Bank.

### Escape phage mutants

To gain further insights into the mechanism of PinR phage inhibition, T2 escape mutants were generated and sequenced. Escape mutant populations were generated by sequentially propagating T2 phage on the same *E. coli* host so that only the phage undergoes mutation. Overnight cultures of BW25113/pCA24N-*pinR* were cultured in LB Cm30 to the exponential phage (turbidity at 600 nm ∼ 0.5), and after 2 h induction with IPTG, T2 phage was added, and cells were incubated for 16 h in 25 mL in 250 mL shake flasks at 37°C while shaking at 250 rpm. After each batch culture, phages were separated from bacterial debris via centrifugation (10 min at 5,000 rpm), and the supernatants were filtered (0.22 µm filter). The filtered phage lysate was used to infect a fresh culture of BW25113/pCA24N-*pinR* cells for 8 cycles. To determine the extent of T2 phage resistance to PinR-activated inhibition after each cycle, the susceptibility of exponentially-growing (turbidity ∼ 0.5 at 600 nm), PinR-producing cells in 96-well plates (after 2 h induction with IPTG) was monitored in the presence of phage isolated from each batch culture for 12 h by scanning every 10 min with a Tecan Sunrise UV-Vis spectrometer.

Phages from the 8^th^ sequential batch culture were sequenced by extracting phage DNA via the Norgen Biotek Phage DNA Isolation Kit. The presence of T2 phage DNA was confirmed using primers gp28_Fow and gp28_Rev for *gp028* (**Table S2**), and the absence of chromosomal *E. coli* DNA was confirmed using the *rrsG* gene (rrsG_Fow and rrsG_Rev, **Table S2**).

The escape T2 phage DNA was sequenced using Next Generation Sequencing with Illumina 2x150 bp configuration by Genewiz. Sequenced data was uploaded on Galaxy web platform (usegalaxy.org) to analyze the data (38). Specifically, quality trimming was performed on the raw reads using SICKLE (version 1.33.2). Phage genomes were assembled using SPAdes genome assembly tool (3.15.5+galaxy2). Kracken2 (2.1.3+galaxy1) was used to identify contigs that are not part of Tequatrovirus classification to remove host chromosomal DNA contamination from assembled genome. Prokka (1.14.6+galaxy1) was used to annotate assembled genome. GenBank files of sequenced phages were deposited under the accession codes listed in **Table S3**.

### Statistical analysis

Statistical analysis was performed using GraphPad Prism. All data presented are the mean ± one standard deviation, and a Student’s T-test was used to evaluate the difference between data sets (probability values (p) < 0.01 were considered significant).

## RESULTS

### PinR inhibits T2 phage infection

After finding the *pinR* deletion makes *E. coli* more sensitive to T2 infection (28), we sought to explore the extent uncharacterized PinR inhibits T2 infection. Exploring structural insights using AlphaFold (39), we found PinR has all the structural features of a small serine recombinase (40), including a DNA binding site, arm region, and catalytic domain (**Fig. 1A**).

**Fig. 1.**
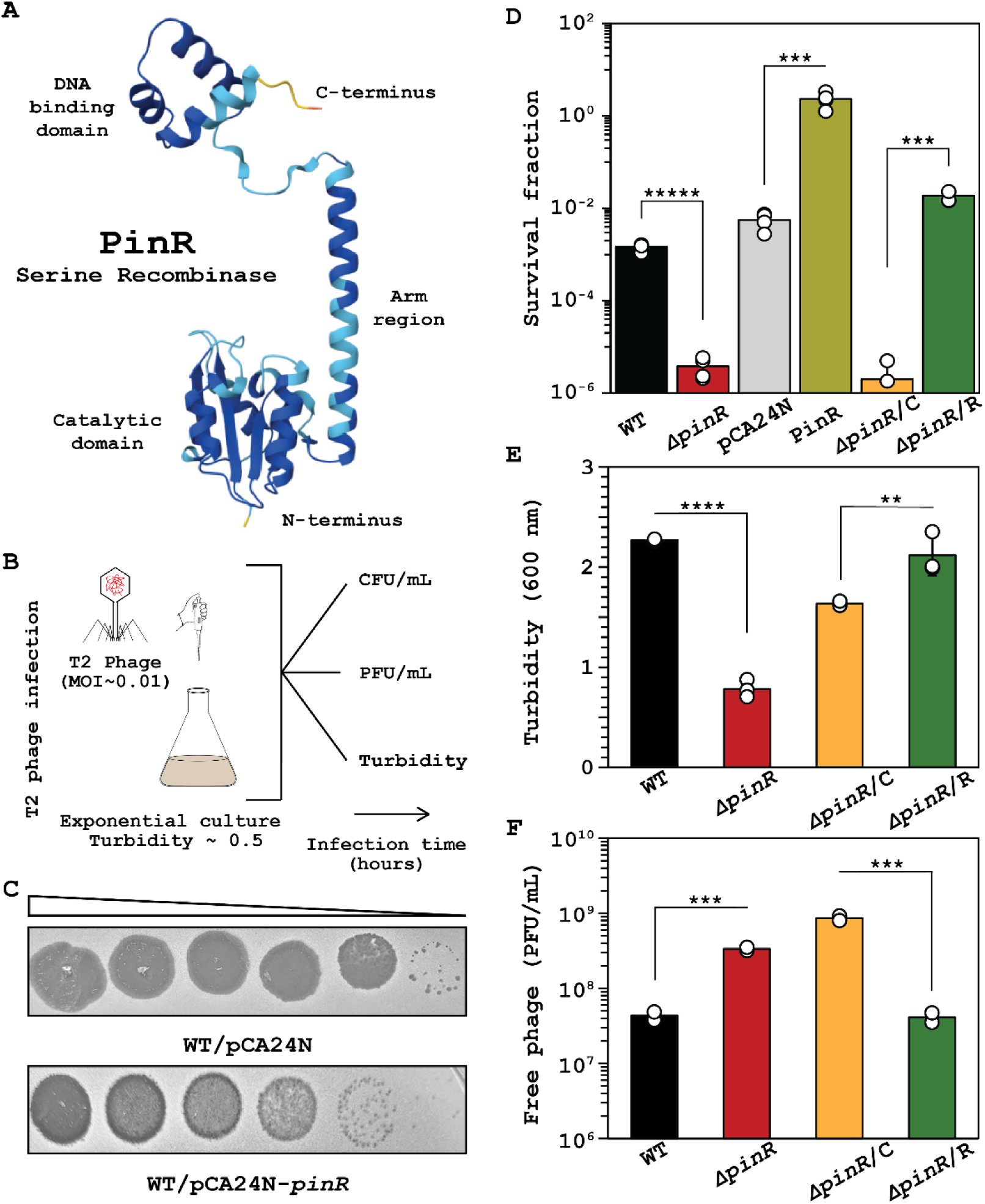
PinR recombinase defends against T2 infection. **A.** PinR is a putative small serine recombinase based on its predicted structure. **B.** Experimental setup for investigating T2 phage inhibition (0.01 MOI) via plaque, CFU, turbidity, and PFU assays. **C.** Plaque assay with cells producing PinR from BW25113/pCA24N-*pinR*. Ten-fold serial dilution of T2 phage. The images shown are for one representative of two independent cultures. **D.** Cell survival after contact with T2 phage (0.01 MOI) for 1 h (full temporal data in **Fig. S1A**). Turbidity (**E**, full temporal data **Fig. S1B**) and T2 free phage (**F,** full temporal data **Fig. S1C**) with T2 (0.01 MOI) after 4 h. Bars and error bars are the mean and standard deviation of four independent cultures. Dots are individual data points. ** p<0.05, *** p<0.005, **** p<0.0005, ***** p< 0.00005. Note: WT is *E. coli* BW25113, pCA24N is WT/pCA24N, PinR is WT/pCA24N-*pinR*, Δ*pinR*/C is *ΔpinR*/pCA24N, and *ΔpinR*/R is *ΔpinR*/pCA24N-*pinR*. IPTG was used at 1 mM for producing PinR from pCA24N-*pinR* in **C**, **D**, **E**, and **F**.

Next, we investigated directly the role of PinR on T2 phage inhibition by assaying plaque formation, host survival, temporal cell viability, temporal cell turbidity, and temporal phage production (experimental design shown in **Fig. 1B**). We found producing PinR from a plasmid reduces T2 plaque formation by ∼10-fold compared to the empty plasmid (**Fig. 1C**). Corroborating this result, in the absence of PinR, (Δ*pinR*), there was a 400-fold decrease in cell survival after 1 hour of infection with T2 phage treatment relative to the wild-type (**Fig. 1D**). We used 0.01 MOI for all experiments to avoid premature cell population collapse. This phenotype was completely complemented since production of PinR (Δ*pinR/*pCA24N-*pinR*) increases cell survival (400-fold increase) relative to the empty plasmid (**Fig. 1D**).

Corroborating the increase in plaques and reduced survival for the Δ*pinR* mutant, temporal cell viability over 8 h without *pinR* in the presence of T2 is significantly reduced (100-fold, **Fig. S1A**). Without PinR, there is also a significant decrease in turbidity during T2 infection (**Fig. 1E**; **Fig. S1B**); this phenotype was also complemented since producing PinR in the *pinR* mutant leads to no observed decrease in turbidity during T2 treatment (**Fig. 1E**; **Fig. S1B**). Corroborating this result, temporal free phage production in the *pinR* mutant is consistently 10-fold higher compared to wild-type strain over 8 h (**Fig. 1F**; **Fig. S1C**), and producing PinR in the *pinR* mutant led to a 20-fold decrease in free phage, complementing the phenotype. Together, these five sets of phage infection studies demonstrate conclusively the activation of PinR inhibits T2 infection.

### PinR inhibits T2 phage infection by inverting the P segment of *e14* cryptic prophage

To determine the mechanism of PinR-related T2 phage defense, we hypothesized that since producing PinR during T2 infection inhibited T2 phage infection (**Fig. 1CDEF**), producing PinR, a putative serine recombinase, must lead to genetic changes that may be discerned by sequencing surviving/growing cells from inside T2 plaques formed on a lawn of PinR-producing cells (**Fig. 2A**). Therefore, whole genome sequencing was performed for surviving cells from two separate colonies in T2 plaques. Strikingly, we found an inversion of a 1,797 bp of cryptic prophage *e14* known as the P segment (27) for both sets of surviving cells (**Fig. 2A**; **Table S4**). As expected for a small serine recombinase, this 1,797 bp fragment is flanked by the two palindromes required for DNA inversion (41) (5’-TTGGTTTGGGAGAAGG-3’ and 5’-CCTTCTCCCAAACCAA-3’) (**Fig. S2**). This inversion occurs within the genes *stfP^+^* and *stfE^+^* in *e14* (**Fig. 2A**; **Fig. S2**); note these two genes are in the opposite orientation, so the inversion, which occurs within the coding region of both, generates two spliced complete genes that we call *stfP2^+^* (begins with *stfP^+^* and ends with *stfE^+^)* and *stfE2^+^* (begins with *stfE^+^* and ends with *stfP^+^*), encoding spliced proteins StfP2 and StfE2 (**Fig. S3**). Hence, cells that survive T2 infection have an inversion of the 1,797 bp P segment in *e14*.

**Fig. 2.**
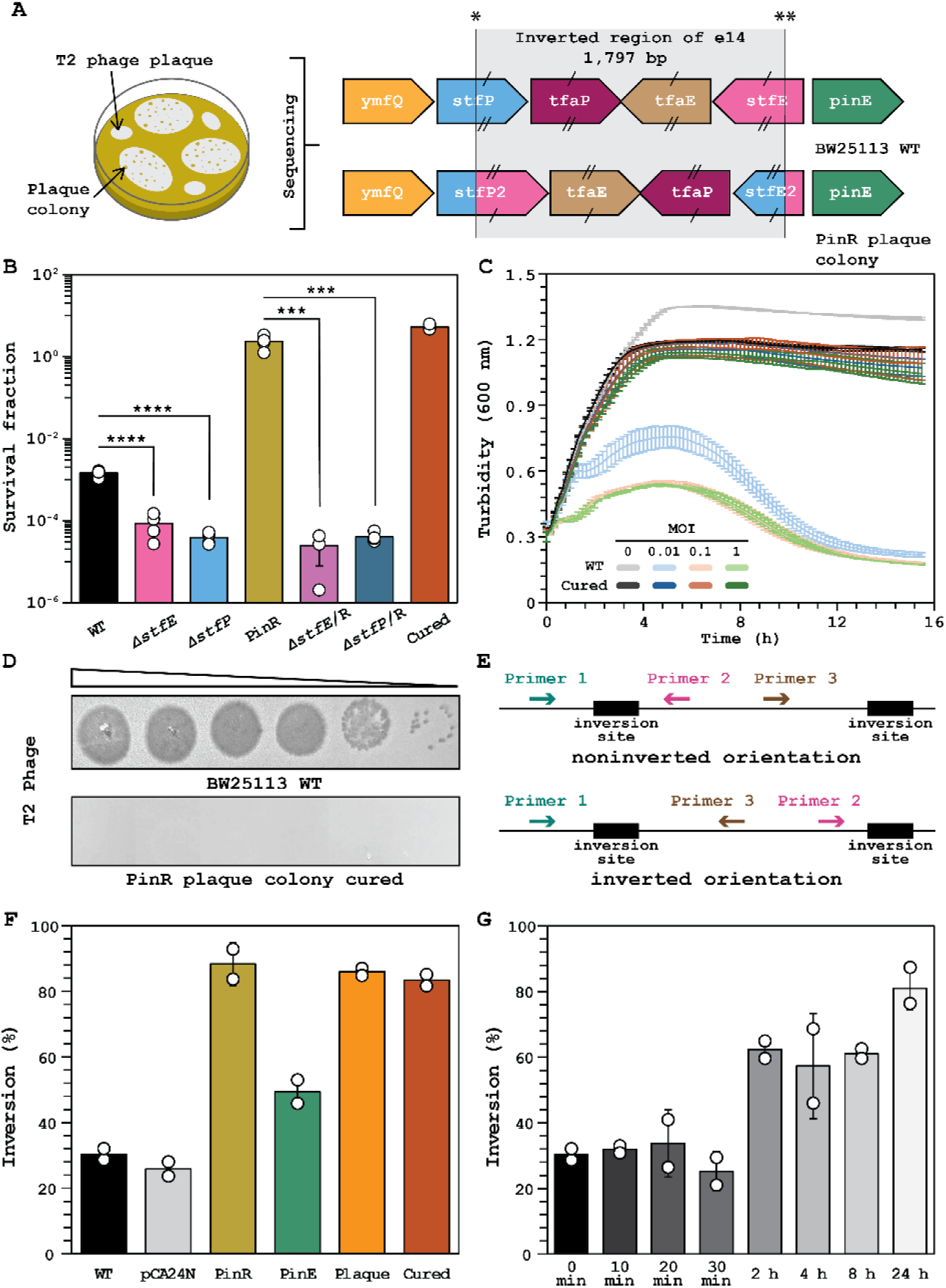
PinR inhibits T2 phage infection by inverting the *e14* P segment. **A.** LHS: schematic of the experiment for isolating colonies within T2 plaques. RHS: sequencing of the plaque survivors revealed the inverted 1,797 bp region (P segment) of *e14* cryptic prophage, which produces spliced genes *stfP2^+^* and *stfE2^+^*. **B.** Cell survival after contact with T2 phage (0.01 MOI) for 1 h. **C.** Turbidity with T2 (0, 0.01, 0.1, and 1 MOI). Data points are the mean and standard deviation of three independent cultures. **D.** Plaque assay. Ten-fold serial dilution of T2 phage WT. Images shown are for one representative of two independent cultures. **E.** qPCR experiment schematic for determining the extent of inversion. **F.** Inversion of DNA isolated from overnight cultures. **G.** Inversion during T2 (0.01 MOI) phage infection. Bars and error bars are the mean and standard deviation of four independent cultures. *** p<0.005, **** p<0.0005. Dots are individual data points. Extent of Inversion determined using primer pair PinR_1 – PinR_2 (noninverted) and PinR_1 – PinR_3 (inverted) (**Table S2**) Note: WT is *E. coli* BW25113, pCA24N is WT/pCA24N, PinR is WT/pCA24N-*pinR*, PinE is WT/pCA24N-*pinE*, Plaque is PinR plaque colony, and Cured is cells derived from the PinR plaque colony cured of pCA24N-*pinR*. Inversion occurs at the palindromes sites * (5’-TTGGTTTGGGAGAAGG-3’) and ** (5’-CCTTCTCCCAAACCAA-3’). IPTG was used at 1 mM for producing PinR from pCA24N-*pinR* in **B** and **F**.

To verify that the sequenced cells from colonies that formed in the T2 plaques are resistant to T2 phage, host survival, temporal turbidity, and phage production were assayed. We found the cured strain (plaque colony cured of the pCA24N-*pinR* vector) had a 3,600-fold increase in cell survival relative to wild-type (**Fig. 2B**). Corroborating this result, the cured strain showed no collapse in growth in the presence of T2 even at 1 MOI (**Fig. 2C**). Strikingly, unlike the wild-type, no T2 plaques were seen at any T2 phage concentration for the cured strain (**Fig. 2D**). Hence, the cells that survived with the DNA inversion in *e14* cryptic prophage are resistant to T2 infection. Corroborating the importance of the inversion, we found preventing inversion either by deleting *stfP* or deleting *stfE*, since each mutation removes one of the palindromes, there was a 40- and 20-fold decrease in cell survival respectively after 1 hour of infection with T2 phage relative to the wild-type (**Fig. 2B**). Hence, the inversion is necessary for phage defense. Similarly, PinR production in the *stfP* and *stfE* mutants (*ΔstfP*/pCA24N-*pinR* and *ΔstfE*/pCA24N-*pinR*) had no impact relative to the *stfP* and *stfE* mutants alone, but deleting *stfP* and *stfE* with PinR heterologous expression led to a 60,000- and 90,000-fold decrease in survival respectively relative to PinR expression in wild-type (BW25113/pCA24N-*pinR*) (**Fig. 2B**).

To confirm PinR inverts the *e14* DNA and to quantify the extent of inversion, qPCR was performed using a three-primer design shown in **Fig. 2E**. We found the inversion was 30 ± 2% in the wild-type and that the empty plasmid pCA24N does not significantly affect inversion (26 ± 3%) (**Fig. 2F**); these inversion rates were corroborated using PCR (**Fig. S4**). In contrast, producing PinR leads to *e14* DNA inversion (88 ± 6%) (**Fig. 2F; Fig. S4**). PinE, a small serine recombinase that is part of cryptic prophage *e14* that inverts this same segment of DNA and has 38% identity with PinR, led to 49 ± 5% inversion (**Fig. 2F; Fig. S4**); like PinR, the physiological role of PinE is unknown (27). Moreover, sequenced cells that survived T2 infection that were isolated from plaques had 86 ± 2% inversion. After curing the sequenced plaque strain of the pCA24N-*pinR* plasmid, the inversion percent is 83 ± 3%, indicating that the inversion is stable, which is expected since Ser recombinases cause irreversible recombinations (18). Hence, PinR is active and inverts *e14* DNA more than PinE. Corroborating these results, we found P segment inversions in 99 *E. coli* genome sequences deposited in the NCBI Gene bank; for example, *E. coli* MG1655 has been sequenced in the inverted and noninverted orientations.

To determine whether the *e14* inversion has physiological relevance for surviving T2 phage infection, temporal *e14* inversion was measured for the wild-type strain during T2 infection. We found the wild-type inverts its *e14* DNA significantly from 30% to 62 ± 4% within 4 h and inversion reaches a maximum after 24 h (81 ± 7%) (**Fig. 2G**). Hence, the P segment of *e14* is inverted in the wild-type strain with native levels of PinR (i.e., no overproduction) during T2 phage infection, and this inversion constitutes a significant phage defense system.

### PinR mediated T2 phage inhibition decreases phage adsorption

So far, we have shown that PinR is active and rapidly inverts the *E. coli e14* P segment to prevent T2 infection, leading to complete resistance. Since many prophages defend their host from rival phages (42) and since preventing attachment is the most prevalent phage defense (43-45), we explored whether P segment inversion by PinR affects T2 adsorption. Remarkably, we found production of PinR significantly decreases T2 phage adsorption with 0% phage adsorption at 8 min and 20 ± 10% adsorption at 16 min (**Fig. 3A**). Corroborating these results, the cured strain had 4 ± 2% and 3 ± 3% adsorption at 8 min and 16 minutes respectively (**Fig. 3A**). In comparison, *E. coli* wild-type strain and empty plasmid (BW25113/pCA24N) had 100% adsorption of T2 phage after 8 min. Hence, PinR reduces T2 phage adsorption.

**Fig. 3.**
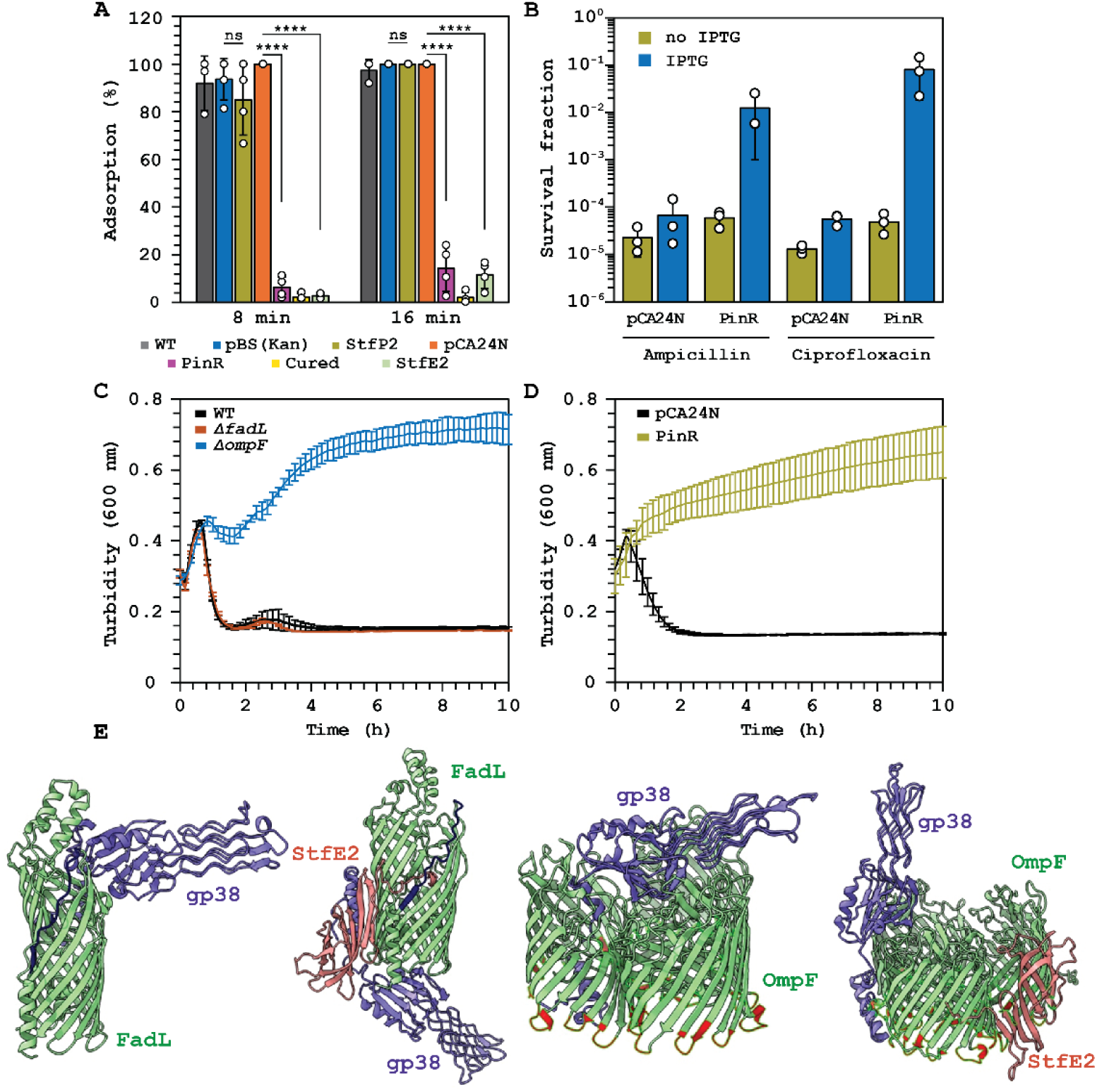
StfE2 inhibits T2 adsorption and PinR reduces porin-related antibiotic import. **A.** T2 phage adsorption on *E. coli* strains. *** p<0.005, **** p<0.0005. **B.** Fold change in cell survival after 3 h treatment with ampicillin (100 µg/mL, 10X MIC) and ciprofloxacin (5 µg/mL, 10X MIC). Bars and error bars are the mean and standard deviation of four independent cultures. Dots are individual data points. Temporal turbidity (600 nm) during ampicillin (**C,** 20 µg/mL) and (**D,** 50 µg/mL,) treatment for exponentially-growing cells. Data points are the mean and standard deviation of four independent cultures. **E.** StfE2 blocks T2 Gp38 adhesion protein on FadL and OmpF. Molecular docking analysis shows interactions of StfE2 with T2 phage adhesion protein Gp38 and T2 receptors, FadL and OmpF. WT is *E. coli* BW25113, pCA24N is WT/pCA24N, pBS(Kan) is WT/pBS(Kan), PinR is WT/pCA24N-*pinR*, StfE2 is WT/pCA24N-*stfE2*, StfP2 is WT/pBS(Kan)-*stfP2*, and Cured is cells derived from the PinR plaque colony cured of pCA24N-*pinR*. IPTG was used at 1 mM for producing PinR from pCA24N-*pinR* in **A**, **B**, and **D**.

To determine if the decrease in T2 phage adsorption was due to a decrease in T2 receptor binding, we explored whether the PinR mediated inversion alters the two T2 receptors, FadL (long-chain fatty acid outer membrane channel) and OmpF (outer membrane porin F) (46) by determining whether transport through the pores formed by FadL and OmpF is affected. Since OmpF imports β-lactam antibiotics (47) and quinolones (48), we reasoned that if the PinR inversion affects the T2 receptor OmpF, ampicillin and ciprofloxacin resistance would be increased by PinR. In agreement with our hypothesis, we found cells producing PinR survive both 10 MIC ampicillin treatment (100 µg/mL) and 10 MIC ciprofloxacin treatment (5 µg/mL) 100-fold and 1000-fold, respectively, relative to cells with the empty plasmid (**Fig. 3B**). Hence, PinR reduces the effect of both β-lactam and quinolone antibiotics.

To determine whether the PinR effect was mediated by OmpF or FadL, we tested which pore was primarily responsible for transporting ampicillin and found deleting *ompF* protects cells from ampicillin at 2 MIC (20 µg/mL) relative to wild-type or the Δ*fadL* mutant (**Fig. 3C**). Similarly, producing PinR protects the cell from ampicillin at 5 MIC (50 µg/mL) (**Fig. 3D**). These results suggest indirectly that the PinR phage defense mechanism is tied to blocking at least the OmpF T2 phage receptor.

### StfE2 blocks T2 adsorption

Since the PinR-mediated P segment *e14* inversion affects four genes (*tfaP^+^, tfaE^+^, stfP2^+^,* or *stfE2^+^*) of the inverted P segment (**Fig. 2A**), we sought to determine which proteins encoded by these four genes are responsible for T2 phage inhibition. To avoid polar effects from gene knockouts, we produced each of the four proteins (rather than studying deletions) during T2 infection and found producing the putative tail fiber TfaE (BW25113/pCA24N-*tfaE*) leads to no survival and producing the putative tail fiber TfaP (BW25113/pCA24N-*tfaP*) leads to a 20,000-fold reduction in host survival (**Fig. S5A**). Corroborating these results, producing TfaP and TfaE also reduced turbidity during T2 infection (**Fig. S5B**). Moreover, TfaP and TfaE production causes only a slight decrease in the growth rate (**Fig. S5C**), which would lead to increased cell survival during T2 infection rather than the increased cell death as seen. Hence, these two proteins do not provide phage defense but are instead detrimental to survival. In contrast, producing StfE2 dramatically reduced T2 phage adsorption with 10 ± 10% phage adsorption at 8 min and 20 ± 20% adsorption at 16 min (**Fig. 3A**). In addition, producing StfP2 had no effect on T2 adsorption (**Fig. 3A**). Therefore, PinR inhibits T2 phage infection by inverting the P segment, which leads to StfE2 production and blocking T2 adsorption.

Bioinformatic analyses were then used to explore further how the newly-formed by inversion StfE2 protein may affect T2 adsorption. StfE2 residues 47 to 125 include the C-terminal gp53-like domain that is related to pyocins like the R1 pyocin of the prophage LESB58 of *Pseudomonas aeruginosa* PAO1 and is related to a region of the putative cell-binding protein Gp53 from the myophage AP22 that infects *Acinetobacter baumanii* (49,50). Therefore, we hypothesized that StfE-2 may block the binding of T2 to its receptor; Gp38 is the primary T2 adhesin that binds to receptors to allow for its adsorption (51). Molecular docking analysis of StfE2, OmpF, and Gp38 shows that StfE2 may alter the binding of Gp38 to OmpF (**Fig. 3E**). Molecular docking analysis (**Fig. 3E**) of StfE2, FadL, and Gp38 shows that StfE2 may also alter drastically the interaction of Gp38 with FadL. These results suggest that StfE2 may directly inhibit T2 adsorption to *E. coli*.

### T2 phage escapes PinR inhibition by mutating *gp38*

To glean further insights into the PinR phage inhibition system, T2 phage was evolved while keeping the production of PinR unvarying **Fig. 4A**. Evolved phages from each cycle were tested for their ability to lyse cells producing PinR (**Fig. 4B**). We found that after two rounds of evolution, the PinR phage inhibition system was increasingly less effective with a collapse in turbidity observed at 3 h by round eight. We then sequenced the T2 genome after eight rounds and compared it to the initial T2 phage (round 0). The T2 escape mutant after eight rounds had point mutations in *α-gt* (encoding an alpha-glucosyltransferase) and in *rnlA* (encoding an RNA ligase and tail fiber attachment catalyst gene) (**Table S5**). Critically, there was also a 48 bp deletion in *gp38*, which encodes the tail fiber protein for host specificity; i.e., for recognizing FadL and OmpF, and results in a 16 aa truncation in hyper variable region 3 (**Fig. 4C**). Hence, T2 phage escapes primarily by mutating the gene responsible for phage adsorption which corroborates the changes in adsorption we found upon activating PinR.

**Fig. 4.**
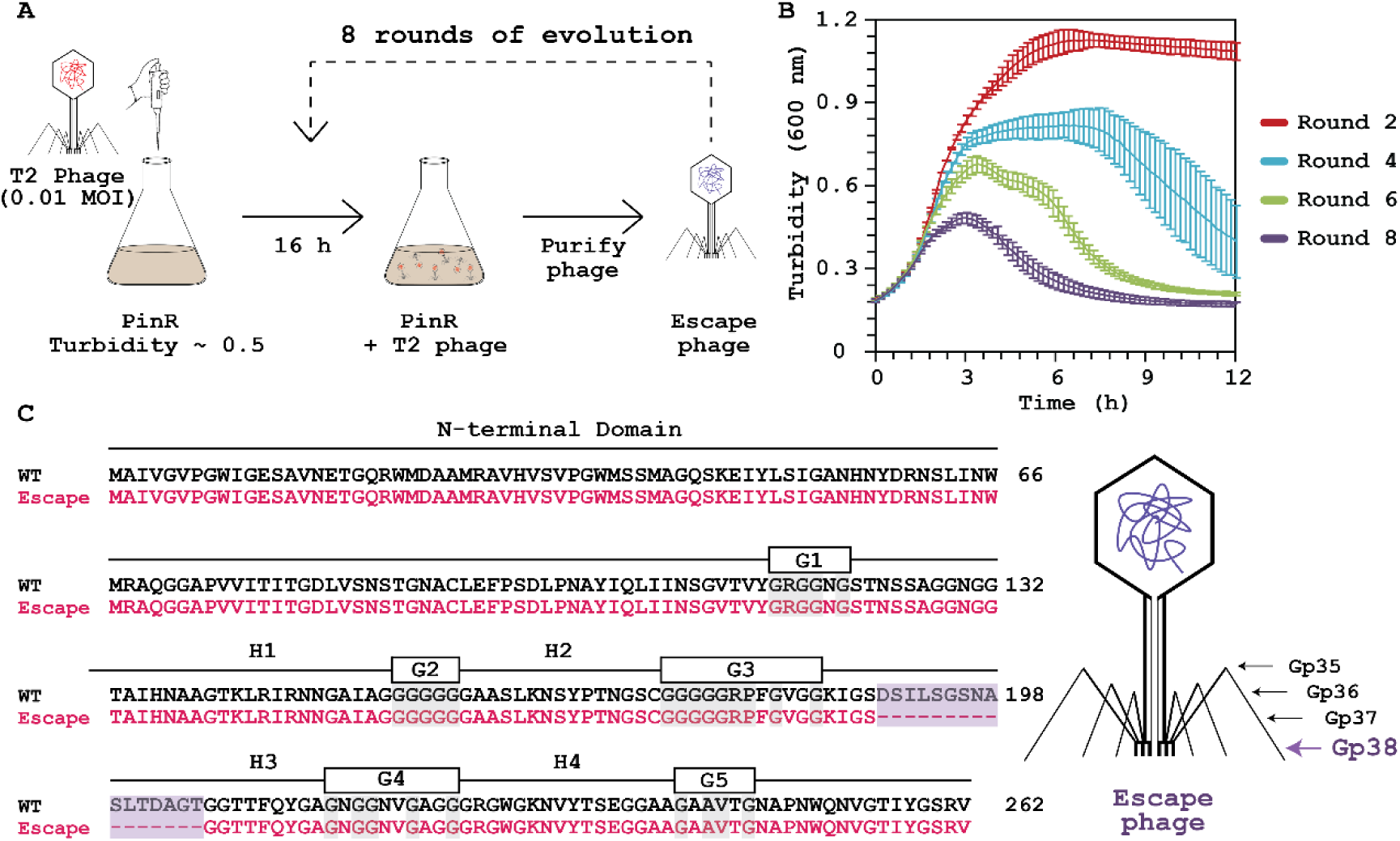
T2 escapes by altering cell adhesion. **A.** Experimental setup for evolving T2 phage against PinR phage defense. T2 phage (0.01 MOI) is added to a fresh, exponentially-growing culture that produces PinR continuously (BW25113/pCA24N-*pinR* with 1 mM IPTG). After 16 h, T2 phage is purified and added to fresh PinR-producing culture that was used for the original round of evolution for 8 evolutions; i.e., T2 phage is allowed to evolve while the host is fixed. **B.** Temporal turbidity as a function of phage infection time for T2 (0.01 MOI) phage from each round of phage evolution. Data points are the mean and standard deviation of three independent cultures. **C.** Comparison of Gp38 adhesin sequences of T2 WT phage (round 0) and T2 escape phage (round 8). Purple highlight indicates the 16 amino acid deletion in Gp38 adhesion gene in the round 8 escape mutant relative to T2 WT phage, and text above the DNA sequences indicate the conserved adhesin’s segments: four hyper variable segments (H1 – H4) and five glycine rice motifs (G1 – G5).

### PinR is widespread

To investigate whether PinR is a widespread phage defense system, we searched UniProt (52) for proteins with more than 90% identity with PinR. The *E. coli* K-12 genome contains PinQ with high similarity (99%) with PinR in *Qin* cryptic prophage. Similarly, PinR is present in 73 genomes of eukaryota (e.g., *Clonorchis sinensis*), bacteria (e.g., *Escherichia* spp. and *Shigella* spp.), and viruses (e.g., *Escherichia* Phages: 2H10, mEp460_ev081, and Tritos). Furthermore, an analysis of proteins with more than 50% identity, found PinR in 623 genomes, including more eukaryota (e.g., Platyhelminthes, Mucoromycete, and Gyrista), bacteria (e..g., *Calditrichota* spp*., Nitrospirota* spp*., Planctomycetota* spp. and *Thermodesulfobacteriota* spp.) (**Table S6**). Corroborating this, recombinases have been found in phage defense islands of multiple bacteria (e.g., *Enterococcus faecalis* (29), *Polymorphum gilvum* (53), *Choloroblium phaeobacteroides* (53), *Desulfovibrio vietnamenis* (54), *E. coli* (55), and *Pseudomonas aeruginosa* (56)). This suggests that PinR phage defense is widespread.

## DISCUSSION

Our results demonstrate that *E. coli* cryptic prophage *rac* encodes an active serine recombinase PinR, that mediatesT2 phage inhibition based on six lines of evidence: PinR (i) increases host survival upon T2 challenge, (ii) reduces T2 phage formation, (iii) inverts the 1,797 bp P segment in cryptic prophage *e14*, (iv) reduces T2 productivity, (v) inverts the P segment of the wild-type strain within 2 h of T2 phage infection, showing the importance of PinR at physiological levels, and (vi) leads to production of inversion-generated StfE2 which decreases T2 adsorption. This phage inhibition system is comparable to that of other active phage defense systems (15,57-61) because PinR-mediated inversion completely inhibits T2 phage infection (**Fig. 2D**). Remarkably, instead of inverting the segment of DNA next to the recombinase, PinR in *rac* cryptic prophage inverts the segment of another cryptic prophage, *e14*; hence, our data provide additional evidence that cryptic prophages play an important role in bacterial physiology (10-12) through their recombinases and that the cell fine tunes its response to phage infection by simultaneously combining resources from distinct cryptic prophages. Similar phenomenon was observed in *Salmonella* LT2, where Fin, a DNA invertase located in prophage Fels-2 controls the inversion of the H segment next to Hin (62). Similarly, PinE inverts the phase determinant region of *Salmonella typhimurium* (63) and G segment of phage Mu (64).

Curing the cell producing PinR isolated from the T2 plaque of its pCA24N-*pinR* plasmid does not cause any significant effect on the inversion present which indicates that the inversion is stable. Corroborating this, the P segment inversion is also observed in nature in numerous sequenced *E. coli* strains. For example, CV601 (NCBI Accession: CP067994) contains the four inversion genes (JJT18_12940 matches *stfE2*, JJT18_12945 matches *tfaE*, JJT18_12950 matches *tfaP*, and JJT18_12955 matches *stfP2*) next to *pinE* that match the sequence of the inverted form of the P segment of the *e14* prophage found in the BW25113 strain isolated here from T2 plaques.

T2 contains six long tail fibers for recognizing primary host receptors and six short tail fibers for recognizing secondary receptors to trigger injection of phage DNA into the host (**Fig. 4C**) (65). Of the long tail fibers, adhesin Gp38 determines the host receptor affinity of T-even phages (66) and for T2, it recognizes FadL and OmpF (46). Gp38 has conserved motifs (i.e., N-terminal domain, four hyper variable segments, five conserved glycine rich motifs and Cfin region) needed to mediate adsorption (67). The T2 phage that escaped the PinR phage defense lacks 16 aa in the hyper variable region 3 (H3 segment) of Gp38; however, it still contains all the conserved motifs of an Gp38 adhesin. Small point mutations in Gp38 adhesins can change host range function significantly; for example, T2 and SV76 phage Gp38 adhesins differ by 2 amino acids but have two different receptors (T2: OmpF and FadL (46); SV79: OmpF and FhuA (68)), which suggests that the 16 aa deletion of the escape mutant identified in this work would cause a complete change in receptor affinity. Collectively, we have identified a new paradigm for phage inhibition via ubiquitous recombinases (**Fig. 5**).

**Fig. 5.**
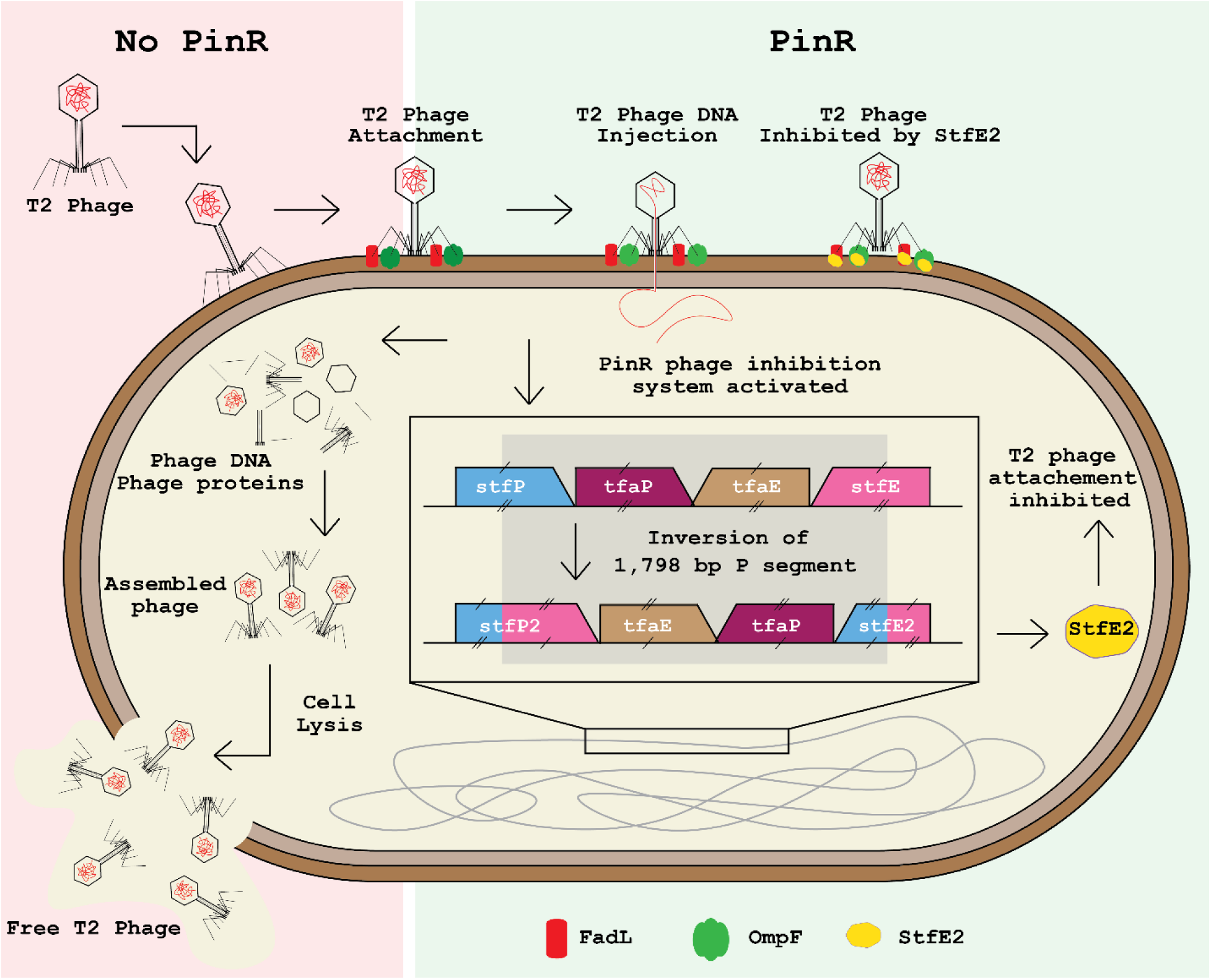
Schematic of the PinR phage inhibition system. **LHS:** Without PinR defense, T2 lyses cells. Free T2 phage lands on the surface of wild-type cell, and the adhesion protein Gp38 recognizes specific T2 phage receptors, OmpF (green) and FadL (red); the T2 lytic cycle then proceeds. **RHS**: PinR-mediated inhibition system. PinR inverts the 1,797 bp P segment of *e14* cryptic prophage within the genes *stfP^+^* and *stfE^+^* and produces two new complete genes *stfP2* (begins with *stfP^+^* and ends with reverse complement of *stfE^+^)* and *stfE2^+^* (begins with *stfE^+^* and ends with the reverse complement *stfP^+^)*. This inversion produces StfE2 which interacts with OmpF and FadL at the same location where Gp38 binds inhibiting T2 phage adsorption.

## DATA AVAILABILITY

Data are available within the body of text and the supplemental information. DNA sequences are deposited in NCBI’s sequence read archive under Bioproject: PRJNA1213105.

## SUPPLEMENTARY DATA STATEMENT

Supplementary Data are available at NAR Online.

## AUTHOR CONTRIBUTIONS STATEMENT

**Joy Kirigo:** investigation, data curation, formal analysis, and writing-original draft preparation. **Daniel Huelgas-Méndez:** software. **Michael J Benedik:** supervision and writing-reviewing and editing. **Rodolfo García-Contreras:** investigation, data curation, supervision, and writing-reviewing and editing. **Thomas K. Wood:** supervision, writing-reviewing and editing, project administration, and funding acquisition.

## Supporting Information

**Table S1.** *Escherichia coli* bacterial strains, plasmids, and phages utilized. Single gene knockout strains in the *E. coli* K-12 BW25113 strain are from the Keio Collection. Plasmids were obtained from the ASKA plasmid library or constructed as indicated. Cm^R^ is chloramphenicol resistance, and Kan^R^ is kanamycin resistance.

| Strains/Phages /Plasmids | Features | Source |
| --- | --- | --- |
| **Strains** |  |  |
| BW25113 | *rrnB3 ΔlacZ4787 hsdR514 Δ(araBAD)567 Δ(rhaBAD)568 rph-1* | (30) |
| BW25113 PinR plaque colony 1 | P segment 1,797 bp *e14* inversion | This study |
| BW25113 PinR plaque colony 2 | P segment 1,797 bp *e14* inversion | This study |
| BW25113 *ΔpinR* | *pinR* Ω Kan<sup>R</sup> | (30) |
| BW25113 *ΔtfaP* | *tfaP* Ω Kan<sup>R</sup> | (30) |
| BW25113 *ΔtfaE* | *tfaE* Ω Kan<sup>R</sup> | (30) |
| BW25113 *ΔstfP* | *stfP* Ω Kan<sup>R</sup> | (30) |
| BW25113 *ΔstfE* | *stfE* Ω Kan<sup>R</sup> | (30) |
| **Plasmids** |  |  |
| pCA24N | Cm<sup>R</sup>; *lacI*<sup>q</sup>, pCA24N | (31) |
| pCA24N-*pinR* | Cm<sup>R</sup>; *lacI*<sup>q</sup>, pCA24N P<sub>T5-lac</sub>::*pinR* | (31) |
| pCA24N-*pinE* | Cm<sup>R</sup>; *lacI*<sup>q</sup>, pCA24N P<sub>T5-lac</sub>::*pinE* | (31) |
| pCA24N-*tfaE* | Cm<sup>R</sup>; *lacI*<sup>q</sup>, pCA24N P<sub>T5-lac</sub>::*tfaE* | (31) |
| pCA24N-*tfaP* | Cm<sup>R</sup>; *lacI*<sup>q</sup>, pCA24N P<sub>T5-lac</sub>::*tfaP* | (31) |
| pCA24N-*stfE2* | Cm<sup>R</sup>; *lacI*<sup>q</sup>, pCA24N P<sub>T5-lac</sub>::*stfE2* | This study |
| pBS(Kan) | Kan<sup>R</sup>; *lacZ*, pBS | [32] |
| pBS(Kan)-*stfP2* | Kan<sup>R</sup>; *lacZ*, pBS P<sub>lac</sub>::*stfP2* | This study |
| **Phages** |  |  |
| T2 |  | TKW stock |

**Table S2.**
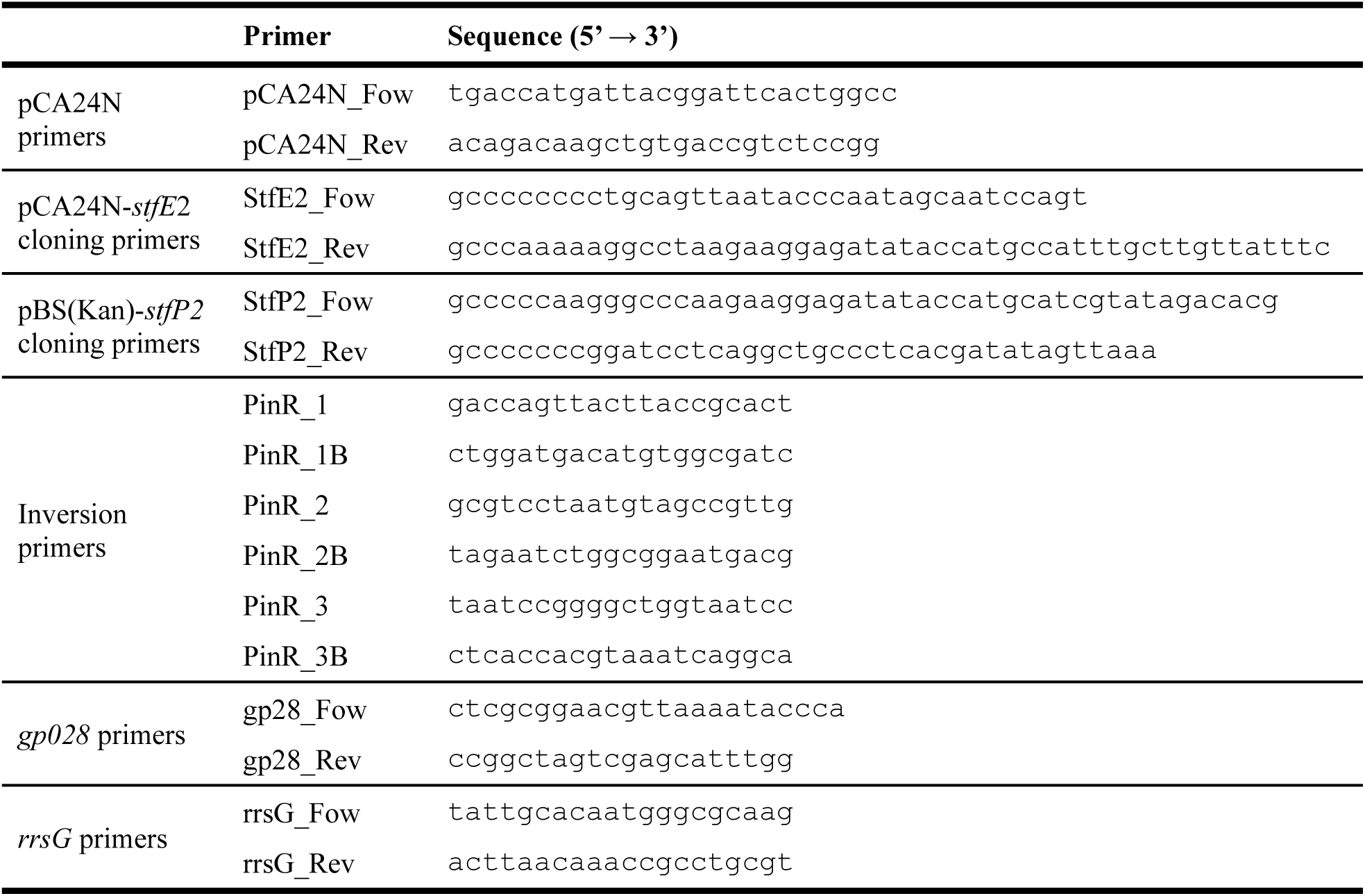
Primers utilized.

**Table S3.** Accession numbers. DNA sequences deposited in NCBI (Bioproject: PRJNA1213105) for the BW25113/pCA24N-*pinR* strains isolated from T2 phage plaques and for the evolved T2 escape mutant. WT is BW25113.

| Sample | Colony | Sample Name | Accession |
| --- | --- | --- | --- |
| **PinR plaque colony** | C1 | WT/pCA24N-*pinR* C1 | SUB15020574 |
|  | C2 | WT/pCA24N-*pinR* C2 | SUB15020780 |
| **T2 phage** | WT | *E. coli* phage T2 | SUB15022426 |
|  | Escape phage | Escape *E. coli* phage T2 | SUB15022439 |

**Table S4.** T2 infection leads to inversion of the *e14* P segment. Mutations in two sequenced BW25113/pCA24N-*pinR* colonies isolated from T2 phage plaques. PinR was produced by overnight induction with 1 mM IPTG then used to produce two-layer TA plates.

| Sample | Results | Genes |
| --- | --- | --- |
| **PinR Plaque 1** | Inversion of 1,797 bp of cryptic prophage *e14* | *stfP* (phage tail protein)<br>*tfaP* (putative tail fiber assembly protein)<br>*tfaE* (putative tail fiber assembly protein)<br>*stfE* (phage tail protein) |
| **PinR Plaque 2** | Inversion of 1,797 bp of cryptic prophage *e14* | *stfP* (phage tail protein)<br>*tfaP* (putative tail fiber assembly protein)<br>*tfaE* (putative tail fiber assembly protein)<br>*stfE* (phage tail protein) |

**Table S5.**
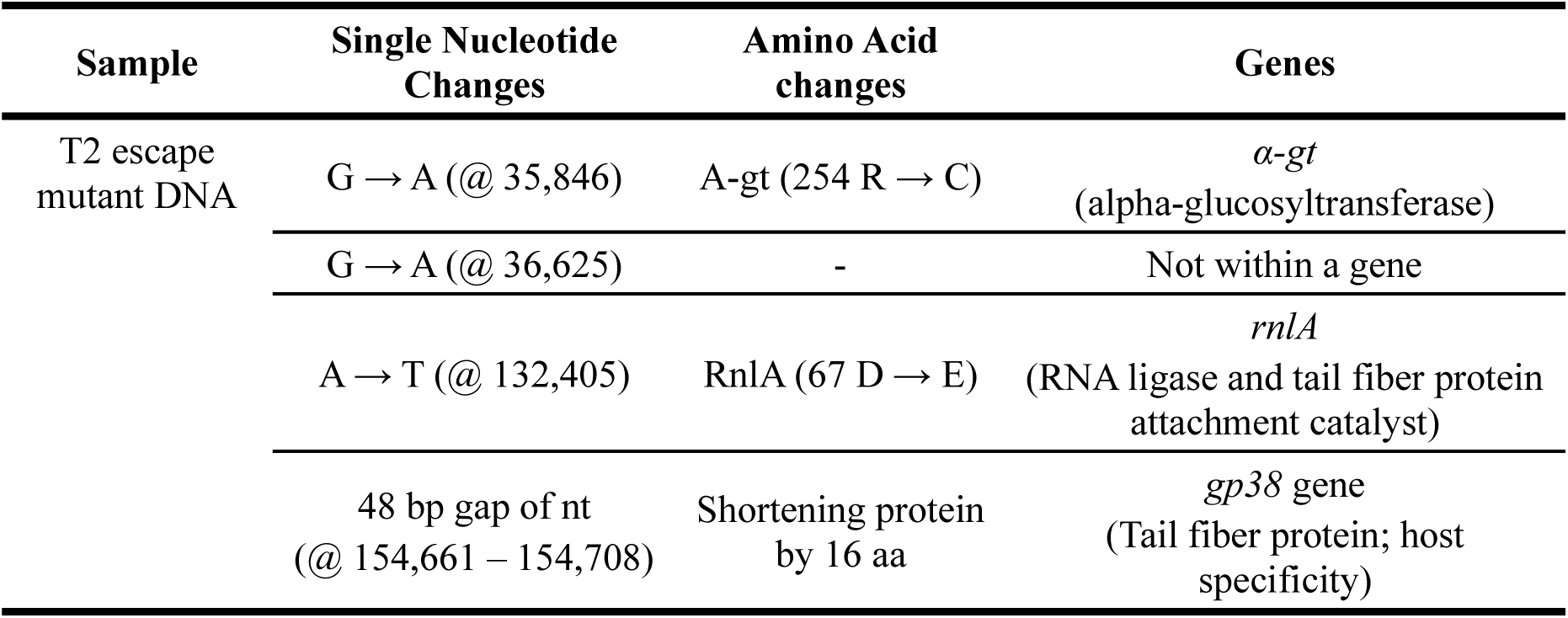
T2 escape mutants. Mutations in T2 phages that escape 8 rounds of sequential contact with BW25113 producing PinR (BW25113/pCA24N-*pinR*) with 1 mM IPTG.

| Sample | Single Nucleotide Changes | Amino Acid changes | Genes |
| --- | --- | --- | --- |
| T2 escape mutant DNA | G → A (@ 35,846) | A-gt (254 R → C) | *α-gt*<br>(alpha-glucosyltransferase) |
|  | G → A (@ 36,625) | - | Not within a gene |
|  | A → T (@ 132,405) | RnlA (67 D → E) | *rnlA*<br>(RNA ligase and tail fiber protein attachment catalyst) |
|  | 48 bp gap of nt<br>(@ 154,661 – 154,708) | Shortening protein<br>by 16 aa | *gp38* gene<br>(Tail fiber protein; host specificity) |

**Table S6.** PinR is widespread. Summary of the kingdoms, phyla, and domains that contain a recombinase with >50% similarity to PinR using UniPort analysis. link:https://www.uniprot.org/uniprotkb?query=uniref_cluster_50:UniRef50_P0ADI0

| Domain | Phylum | Kingdom |
| --- | --- | --- |
| Phages | *E. coli* phages | - |
| Bacteria | *Pseudomonadota (Proteobacteria)* | - |
|  | *Calditrichota* | - |
|  | *Nitrospirota* | - |
|  | *Planctomycetota* | - |
|  | *Thermodesulfobacteriota* | - |
|  | *Nitrospirota* | - |
| Eukaryota | *Platyhelminthes* | Parasitic worm |
|  | *Mucoromycota* | Fungi |
|  | *Gyrista* | Brown algae |

**Figure S1.**
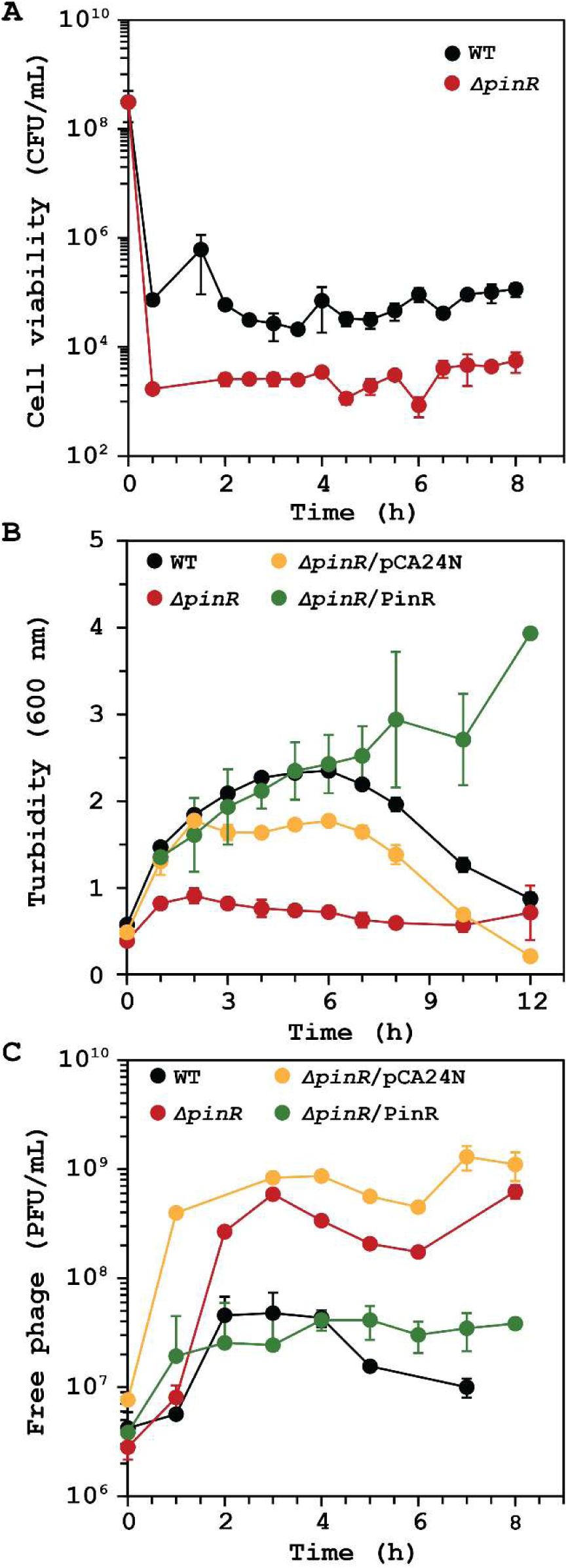
Temporal data showing PinR inhibits T2 infection. Cell viability (**A**), turbidity (**B**), and T2 free phage (**C**) with T2 (0.01 MOI). One-hour data of (**A**) are shown in **Fig 1D**, and four-hour data of (**B** and **C**) are shown in Fig. 1E and Fig. 1F, respectively. Data points are the mean and standard deviation of four independent cultures. Note: WT is *E. coli* BW25113 and pCA24N is WT/pCA24N. IPTG was used at 1 mM for producing PinR from pCA24N-*pinR* in **D**, **E**, and **F**.

**Figure S2.**
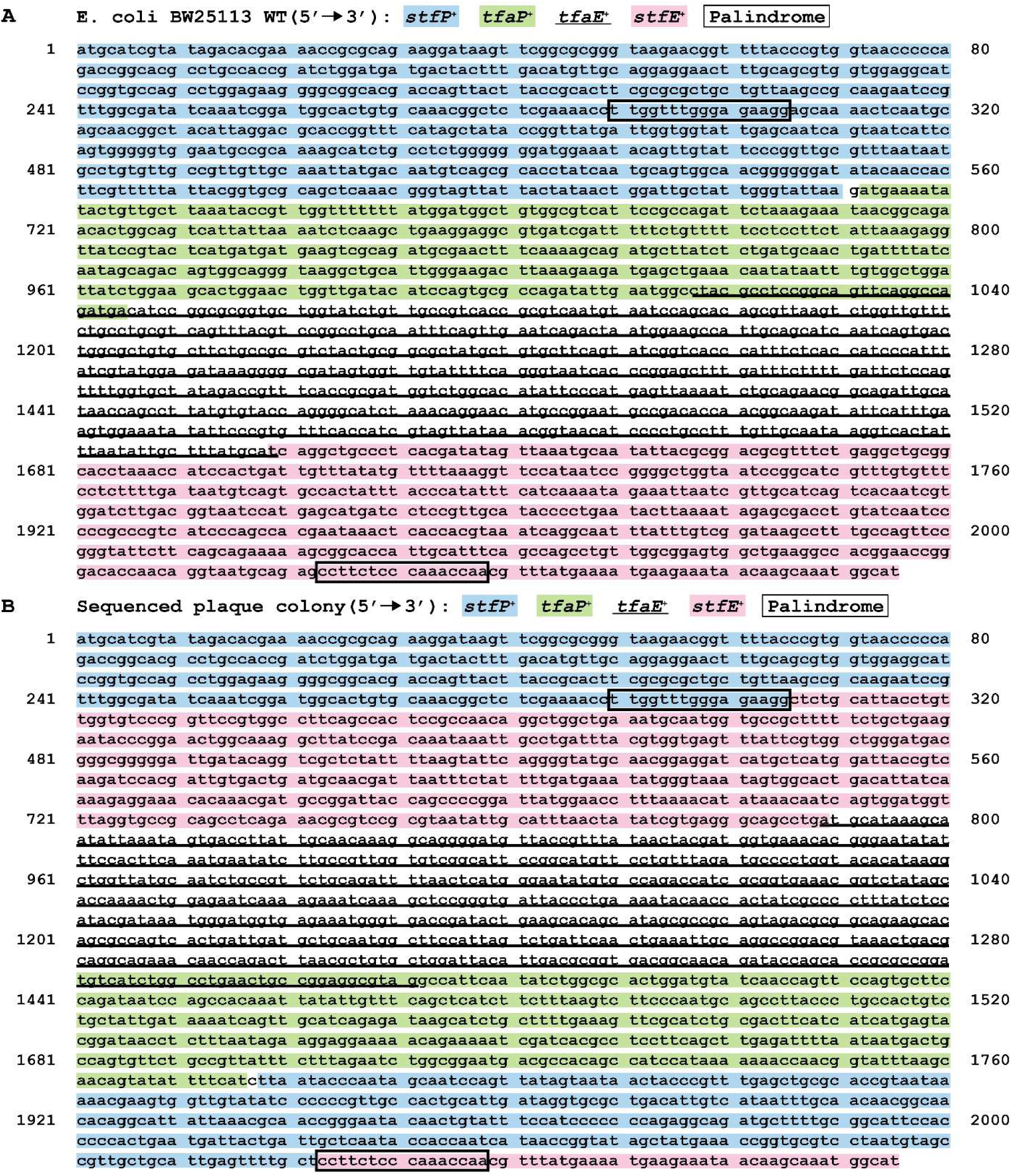
Inverted region of *e14* prophage. Sequence of *stfP^+^, tfaP^+^, tfaE^+^,* and *stfE^+^* genes in *E. coli* BW25113 WT (**A**) and the sequenced PinR-producing plaque colony (**B**), indicating the inverted sequence and palindrome sequences flanking the inverted region (boxed). Light blue: *stfP^+^*; underline: *tfaE^+^*; green: *tfaP^+^*; light pink: *stfE^+^*; box: palindrome sequence flanking the invertible P segment of *e14* prophage.

**Figure S3.**
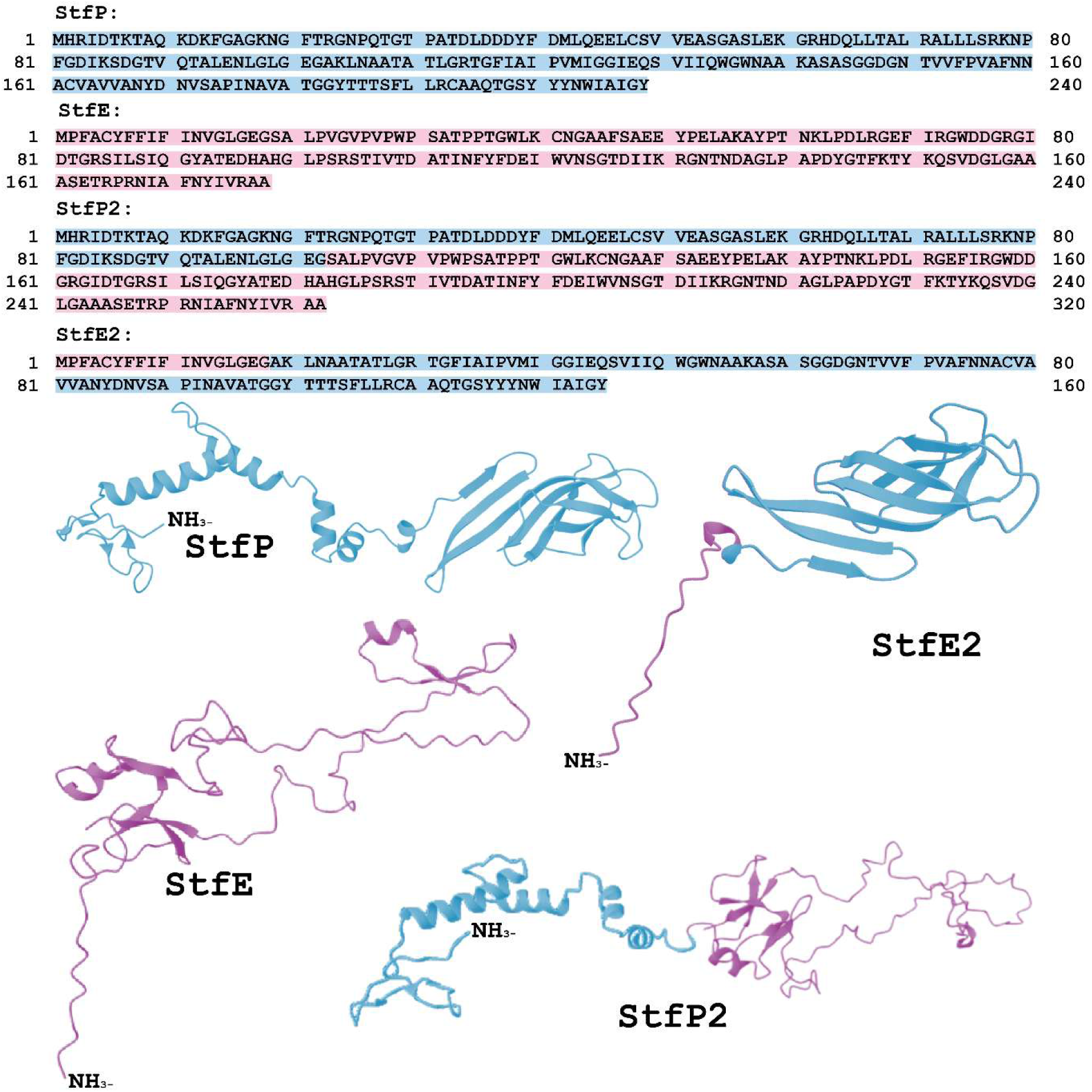
Inversion of P segment forms new spliced proteins StfP2 and StfE2. Primary and tertiary structures of StfP, StfE, StfP2, and SftE2. Light blue indicates StfP residues, and pink indicates StfE residues.

**Figure S4.**
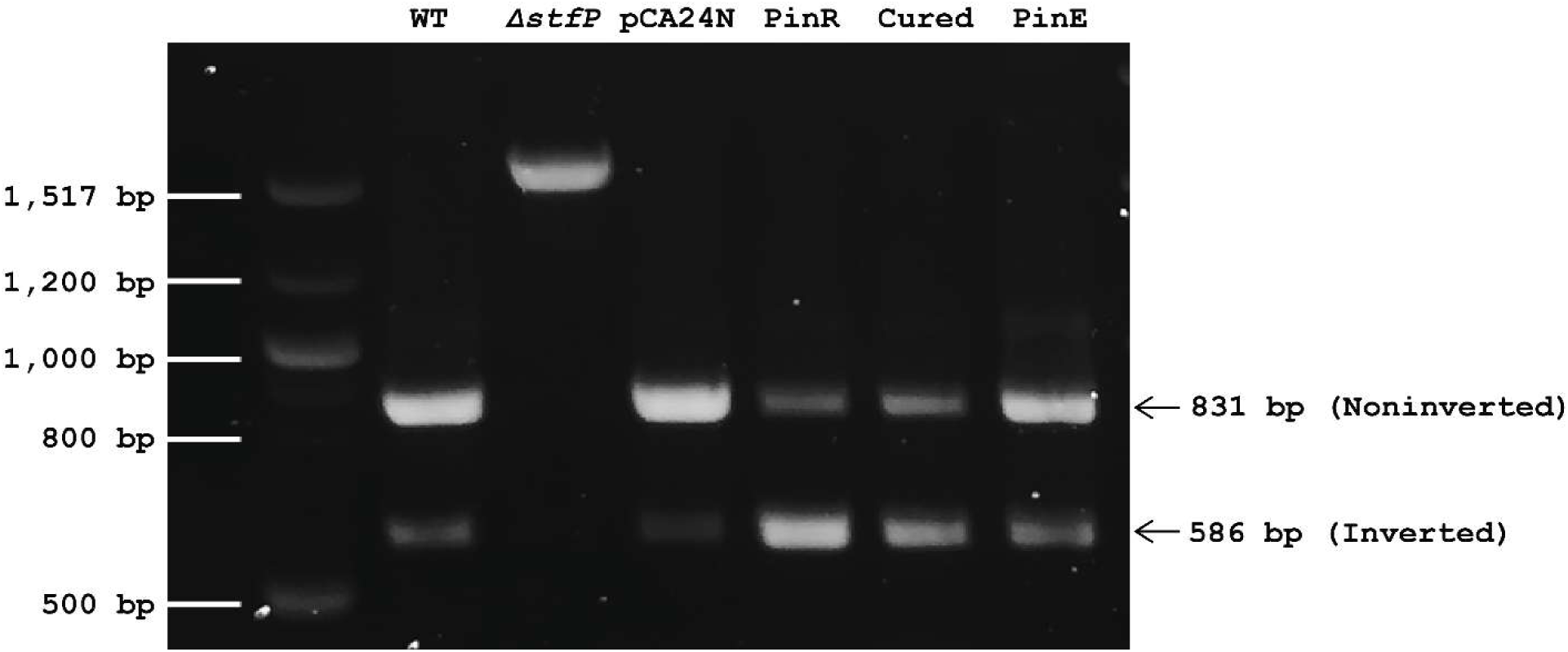
Extent of inversion of the *e14* P segment. Agarose gel electrophoresis of PCR products obtained using primers PinR_1B, PinR_2B, and PinR_3B (**Table S2**). Products are noninverted (831 bp) and inverted (586 bp) except for *ΔstfP* (1,512 bp) due to the Kan^R^ insertion in the KEIO mutant. cDNA samples of WT and pCA24N are predominately noninverted while those of PinR and Cured are inverted. PinE is ∼ 60% noninverted matching results in Fig. 2F. The single band for *ΔstfP* indicates the cDNA sample is 100% noninverted, since the Keio mutant cannot be formed with inverted DNA. Note: WT is *E. coli* BW25113, pCA24N is WT/pCA24N, PinR is WT/pCA24N-*pinR*, Cured is cells derived from the PinR plaque colony cured of pCA24N-*pinR*, and PinE is WT/pCA24N-*pinE.* IPTG was used at 1 mM for producing PinR and PinE.

**Figure S5.**
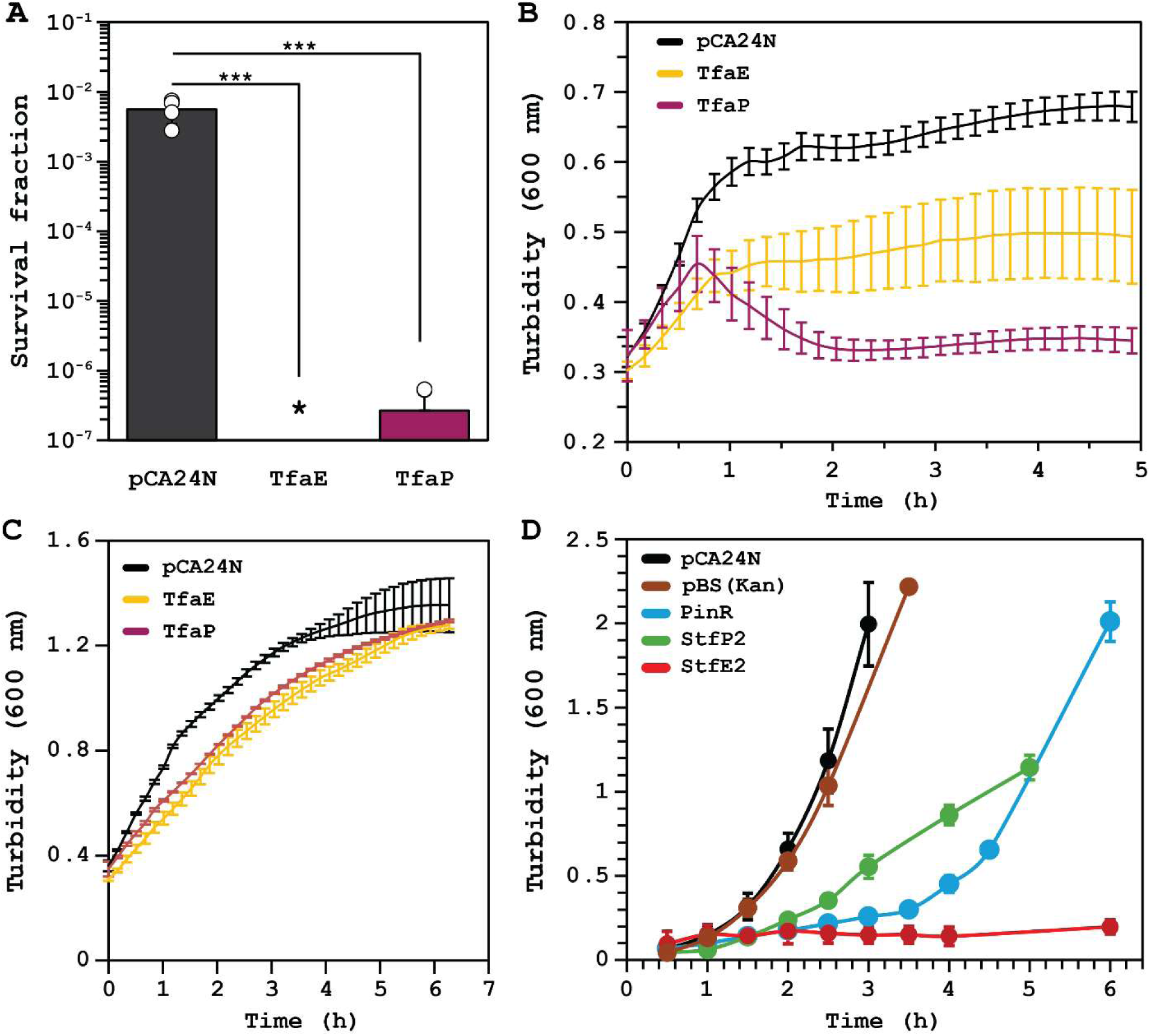
TfaE and TfaP do not inhibit T2 infection and StfE2 is toxic. **A.** Cell survival with T2 phage (0.01 MOI) after 1 hour. *indicates no cell survival. Bars and error bars are the mean and standard deviation of four independent cultures, respectively. Dots are individual data points. data points. *** p<0.005. **B.** Turbidity (600 nm) during T2 phage (0.01 MOI) infection over time. **C.** Growth in 96 wells (turbidity at 600 nm) in LB medium with 1 mM IPTG without T2 phage. **D.** Growth in shake flasks in LB medium with 1 mM IPTG without T2 phage. (**B**-**D**) Data points are the mean and standard deviation of four independent cultures. Abbreviations: WT is BW25113, pCA24N: WT/pCA24N, pBS(Kan): WT/pBS(Kan), TfaE: WT/pCA24N-*tfaE*, TfaP: WT/pCA24N-*tfaP*, StfE2: WT/pCA24N-*stfE2,* and StfP2: WT/pBS(Kan)-*stfP2*.

